# Mechanics limit ecological diversity and promote heterogeneity in confined bacterial communities

**DOI:** 10.1101/2023.12.18.572212

**Authors:** Tianyi Ma, Jeremy Rothschild, Faisal Halabeya, Anton Zilman, Joshua N. Milstein

## Abstract

Multi-species bacterial populations often inhabit confined and densely packed environments where spatial competition determines the ecological diversity of the community. However, the role of mechanical interactions in shaping the ecology is still poorly understood. Here we study a model system consisting of two populations of non-motile *E. coli* bacteria competing within open, monolayer micro-channels. The competitive dynamics is observed to be bi-phasic: after seeding, either one strain rapidly fixates or both strains orient into spatially stratified, stable communities. We find that mechanical interactions with other cells and local spatial constraints influence the resulting community ecology in unexpected ways, severely limiting the overall diversity of the communities while simultaneously allowing for the establishment of stable, heterogeneous populations of bacteria displaying disparate growth rates.

Surprisingly, the populations have a high probability of coexisting even when one strain has a significant growth advantage. A more coccus morphology is shown to provide a selective advantage, but agent based simulations indicate this is due to hydrodynamic and adhesion effects within the microchannel and not from breaking of the nematic ordering. Our observations are qualitatively reproduced by a simple Pólya urn model, which suggests the generality of our findings for confined population dynamics and highlights the importance of early colonization conditions on the resulting diversity and ecology of bacterial communities. These results provide fundamental insights into the determinants of community diversity in dense confined ecosystems where spatial exclusion is central to competition as in organized biofilms or intestinal crypts.

## Introduction

Bacteria commonly exist within dense and complex communities known as biofilms where they compete for limited space and resources [15, 44]. These microbial consortia are generally not well mixed; instead, the biogeography of most biofilms displays hierarchical levels of spatial patterning much like eukaryotic cells within macro-organisms [3,35,39]. This organization, resulting from the complex dynamics of competitive and cooperative interactions within and between species, can lead to emergent properties of functional significance to the consortium as a whole [24, 32].

Compared to the independent populations from which they are composed, bacterial communities are typically more resilient to environmental pressures such as salinity, pH gradients, and UV damage, as well as to treatment with conventional biocides or antibiotics [11, 20]. Unfortunately, drug resistance as well as multiple diseases, such as cystic fibrosis or infective endocarditis, are linked to the organization and function of these microbial populations as well [18, 34].

Individual bacteria within these communities closely interact and may display cooperative social behaviors to benefit neighbouring cells and the population as a whole (e.g., by excreting digestive enzymes or nutrient chelators) or, conversely, employ competitive, antagonistic strategies to reduce the fitness of their neighbours (e.g., by excreting antibiotics, bacteriocins, and other toxins) [31, 50]. Other social interactions include the exchange of genetic material through horizontal gene transfer (HGT) [28] as well as quorum sensing and chemotactic signaling [5]. These chemically-mediated social behaviours can lead to regulation of cell differentiation, coordinate gene expression, and partition tasks among colony members [2, 12].

Biophysical mechanisms arising from mechanical constraints, imposed by both other bacteria and the environment, also play a central role in the population dynamics and spatial patterning of these communities. For instance, rod-shaped cells like *E. coli* tend to align with neighbouring cells and surfaces resulting in an orientational patterning similar to nematic ordering within liquid crystals [8,40,47]. Mechanical interactions between rod-shaped bacteria also lead to the spontaneous generation of heterogeneity and phase-separation between species within model biofilms [16]. Furthermore, mixed-morphotype colonies of *E. coli*, composed of bacillus (rod-shaped) cells and coccobacillus (round) mutants, growing at similar rates, were observed to sort by shape with the coccobacillus cells organizing along the top of the colony and the bacillus cells along the surface and edges [43]. These observations suggest that cell morphology may play a critical, if often overlooked, role in the organization and evolution of a microbial community [6].

Microfluidic devices can be employed to create model, micro-environments in which single bacteria may be observed for extended periods of time. These engineered systems provide a test-bed for ecological theories of microbial population and community dynamics [38] providing insight into bacterial competition within soil-like porous environments [10], the phage susceptibility of microbial communities within structured environments [42], the protection from predation offered by a structured biofilm [51], among other topics. Recently, Koldaeva et al. studied the competitive dynamics of two effectively neutral strains of *E. coli* grown within open-ended microchannels [26].

Within these devices, the rod-like bacteria tended to align into distinct lanes as previously observed [47]. Within each lane, any diversity was lost exponentially fast as the aligned cells rapidly propagated toward and got expelled from the ends of the channel. This behavior can be explained by the collective growth of bacteria in a lane resulting in fixation times that scale logarithmically with channel size rather than algebraically (as in classical ecological models of well mixed populations such as the Moran model [37]). Propagation or invasion between lanes, however, was much slower maintaining a fixated monoculture within individual lanes. However, while Koldaeva et al. focused on the rapid loss of diversity observed within individual established lanes within a microchannel, we are concerned with understanding what influences the overall ecological diversity which can be supported by a microchannel and how this ecology is established starting from a seeded population.

In this paper, to study the factors controlling the establishment and the maintenance of ecological diversity within a confined microenvironment, we use a model bacterial community composed of two distinct, non-motile *E. coli* colonies competing for dominance within open-ended microchannels. We find that a significant fraction of fixation events happen before the formation of ordered and relatively stable lanes. However, the initial conditions and early dynamics strongly influence the eventual diversity among the ordered state within a filled microchannel where species regularly co-exist for extended times. We then consider the effects of cell morphology on competition by tuning the aspect ratio of one or both of the populations by treatment with S-(3,4-Dichlorobenzyl) isothiourea (A22), a compound which targets the MreB protein that controls cell width in *E. coli* [4, 48]. By introducing an A22-resistant mutant, we are able to selectively induce morphological changes in the wild type population while leaving the mutant cells unchanged. From experimental observations of these dual competitive populations, analysed with agent-based simulations and theoretical modeling, we find that mechanical interactions within dense bacterial communities influence the microenvironmental ecology in unexpected ways, greatly restricting the diversity of the microbial community while, simultaneously, maintaining spatially heterogeneous populations where less fit bacteria stably coexist with fitter bacteria.

## Results

Two isogenic populations of non-motile, wild-type *E. coli* (MC1400) cells, made distinguishable by the expression of either eGFP or mCherry, are grown in a series of open-ended microchannels (46x12x1 *μ*m^3^), see Figs. 1A,B. While the cells are isogenic, throughout, we will make reference to the eGFP (green) or mCherry (red) strains. In both strains, expression of the fluorescent proteins is under the control of an

**Figure 1.**
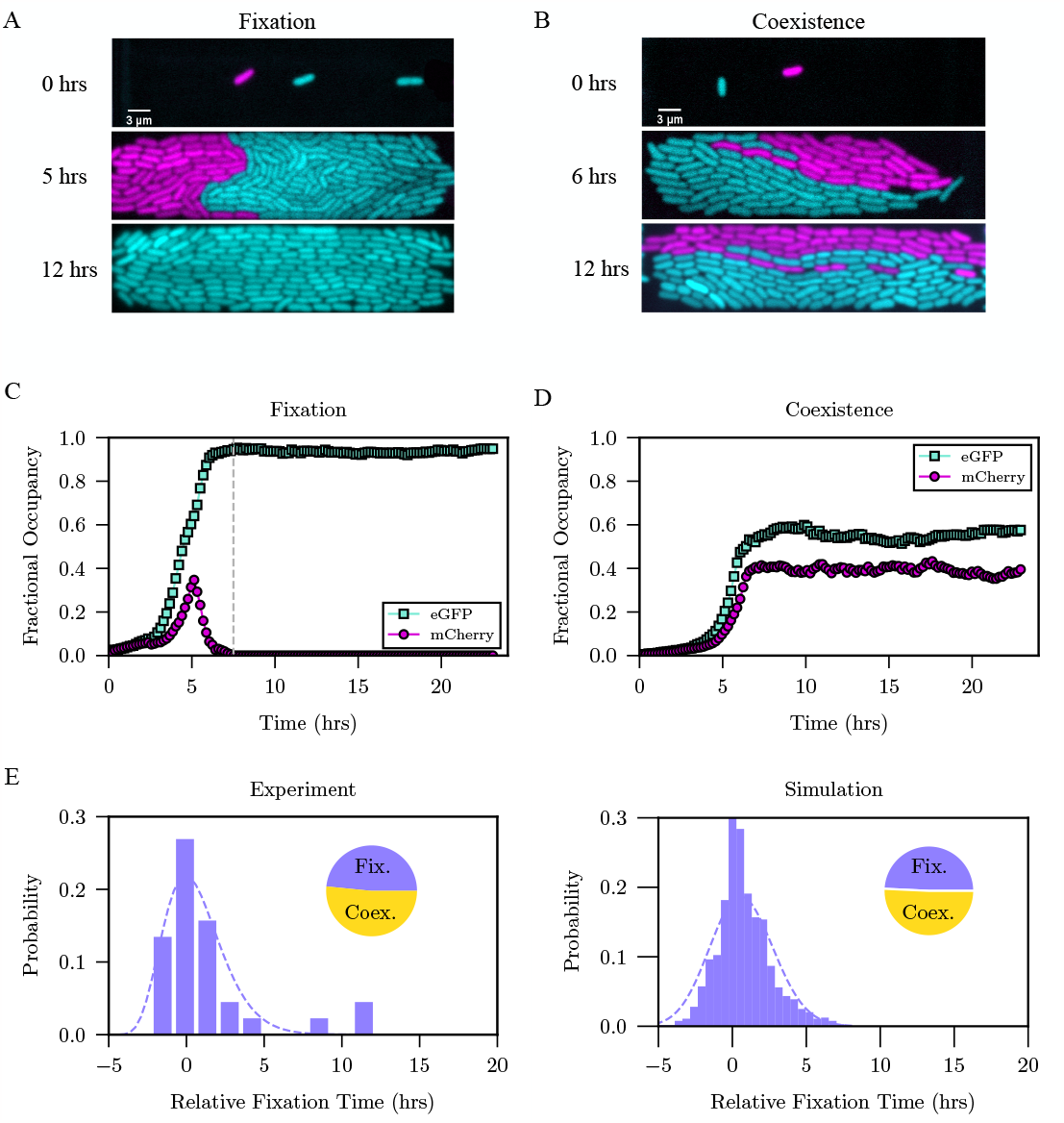
Spatial competition between two *E.coli* strains displays biphasic dynamics resulting in either coexistence or fixation. A,B) Dynamics of competition from low initial seeding within an open monolayer device are captured by time-lapse fluorescence microscopy. Images are shown at *t* = 0, 5 or 6, and 12 hrs. C,D) Total area occupied by each strain normalized to the chamber area as a function of time. Panels A and C provide an example illustrating the observed fixation dynamics, while panels B and D illustrate coexistence dynamics. The grey dashed line in C indicates the point at which the mCherry carrying (red) strain’s occupancy reaches zero, defining the fixation time (7.5 hr). E) Histogram of observed fixation times relative to the average chamber fill time (*t* = 0) fitted with a Gamma distribution (dashed line), for both populations, from competition experiments between roughly neutral *E. coli* strains. The distribution is sharply peaked about the chamber fill time. The inserted pie chart shows an almost equal number of observations of fixation or coexistence (31 fixations out of 64 total experiments). F) Same as in E, but from agent-based simulations (2436 fixations out of 5000 simulations).

L-arabinose inducible P_BAD_ promoter that, as will be discussed, can be used to tune the relative growth rate between strains.

### Establishment of population diversity is biphasic

Both strains are imaged at an initial induction level of 0.01% (w/v) L-arabinose by time-lapse microscopy. Cells are randomly seeded into the microchannels at a low initial abundance (≤ 4 cells/strain) and a similar relative abundance (≤ 3-fold) (see *SI Fig. S10*). A community fixation event is defined as when one population completely displaces the other by pushing it out of the chamber. We find that many fixation events are relatively rapid, occurring around the time it takes dividing cells to fill the entire chamber and before the system orients into nematically ordered, parallel lanes (Figs. 1A,C,E and *Movie S1, S2*). Once the populations reach the ordered state, each lane rapidly fixates as cell divisions drive the populations within each lane toward the open ends of the microchannels. The resulting diversity of the lanes remain locked for a relatively long time because it can only be affected by infrequent inter-lane invasions (Fig. 1D). Based in these observations, we define coexistence in our system to be when both strains are present at the end of the 24-hour observation window as shown in Figs. 1B,D (although, eventually, these systems might fixate as well through successful inter-lane invasions [26, 37]). Unlike the ordered lane-based state of the population, which is amenable to mathematical modeling though discrete stochastic population models [26, 37], the dynamics of the early disordered phase is much more difficult to model. To gain mechanistic insight into our results, we analysed them using minimalistic agent based simulations of bacterial dynamics, which incorporate only the salient features of the physical interactions between the cells (see *SI Section 2*). The simulations (Fig. 1F) reproduced the experimentally observed results, including the biphasic outcomes such as the 50/50 split between fixation and coexistence for cells with rod-like shapes similar to *E. coli*, in good agreement with experiment (Figs. 1E,F).

### Division rate weakly influences the probability of coexistence, but in fixated channels, strongly determines which strain fixates

At either low (0.01% L-arabinose) or high (0.2% L-arabinose) levels of induction of the fluorescent reporters, the eGFP-expressing strain is observed in liquid cultures to grow, on average, slightly faster than the mCherry-expressing strain (see Fig. 2A and *SI Fig. S5)*. This variation in doubling times is thought to result from ROS mediated cytotoxic effects of mCherry [36, 45]. However, when grown at high induction levels within our microfluidic devices, we witness a similar growth advantage for eGFP-expressing cells (on the order of 10%), which is not observed at low induction. Bulk sliding assays, as in [21], similarly display a competitive advantage for the eGFP-expressing population at high induction with cultures seeded at equal cell density, resulting in a significantly larger proportion of green territories (Figs. 2B and *SI Figs. S2, S3*). At low induction, the sliding assays indicate a more neutral competition between strains resulting in a more balanced red/green distribution of territories. Why the induction level appears to affect cells within liquid culture assays differently from those within the more crowded microfluidic and sliding-assays is unclear. Here we simply use the observed imbalance in the rate of cellular growth as a way to adjust the selective advantage of one population over the other within the microfluidics.

**Figure 2.**
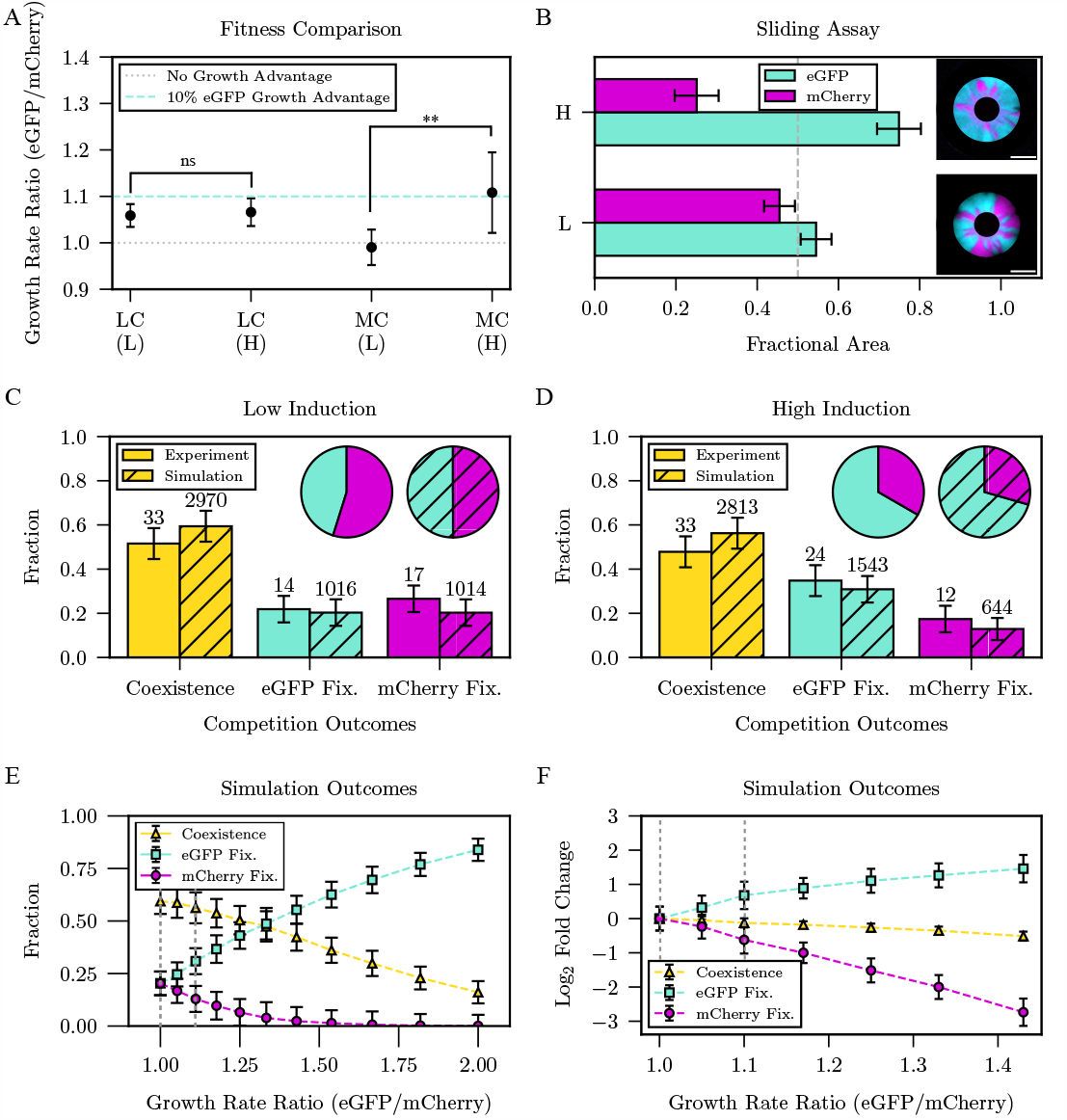
Competition between eGFP and mCherry strains is effectively neutral at low induction while favouring eGFP-expressing cells at high induction. A) Relative difference in doubling times measured in liquid culture (LC) and in microchannels (MC) at low (L) and high (H) induction. Dotted (dashed) lines indicate a 0% (10%) relative growth advantage for eGFP-expressing cells. Error bars are ± the standard deviation of n = 12,12,10,10 (left to right, respectively) replicates. B) Comparison of the fractional area occupied by each strain in sliding assays evaluated at low and high induction. The dashed line at 0.5 indicates balanced competition. Error bars are ± the standard deviation from 3 replicates. Scale bars in representative images are 5 mm. C, D) A comparison of simulations and experiments showing the fraction of microchannels displaying coexistence and fixation at low (C) and high (D) induction. Inset pie charts indicate the fraction of fixations for each strain, observed and predicted, at each induction level. Error bars result from bootstrapping the data. Bootstrapping on the simulated data was subsampled similar to experiments to estimate the expected measurement error (see *SI Section 1*). E) Simulated predictions for the fraction of microchannels displaying coexistence and fixation as a function of the ratio of growth rates (eGFP/mCherry). F) Same as (E) but expressed in terms of log_2_ fold change of the fractions. The two grey dashed lines in (D) and (E) refer to a growth rate advantage of eGFP cells at 0% and 10%, respectively.

For competition experiments within the microchannels, small differences in division time between populations result in significantly more fixation events for the advantaged strain relative to the disadvantaged strain (Fig. 2C,D). Both simulations and experiments show an approximate 2:1 ratio of eGFP to mCherry fixations for a ∼ 10% relative growth advantage by the eGFP strain. However, the probability that one strain fixates over the other is significantly more sensitive to division time than is the likelihood of the cells coexisting. These results are reproduced in simulations where coexistence remains highly probable even for strains with highly disparate growth rates (Fig. 2E,F).

### Well mixed, abundantly seeded populations tend to result in coexistence

We next look at the effect of early colonization conditions, namely, the initial abundance of each strain, on the resulting ecology. In Fig. 3A, we display the experimentally observed competitive outcomes for microchannels randomly seeded at an increasing abundance of each cell type (eGFP & mCherry). All experiments were performed at high induction so that the eGFP expressing strain always maintained a competitive advantage. Because we do not have precise control over the initial seeding, the data were binned into ranges of total abundance and the relative abundance of each strain was restricted (see *SI Section 1, Fig. S10* for seeding distributions). We observed that, for increasingly abundant initial seeding, coexistence becomes increasingly more likely. This behavior is recapitulated by agent-based simulations where we have incorporated a 10% relative growth advantage for the eGFP carrying strain to account for the observed competitive advantage of these cells at high induction. In these simulations, Fig. 3B, the chambers were randomly seeded at an increasing abundance of each strain, but in equal proportions of red/green cells. Our results show that initially well-mixed and abundantly seeded populations tend to coexist at later times highlighting how the initial level of seeding can critically influence the resulting ecology.

**Figure 3.**
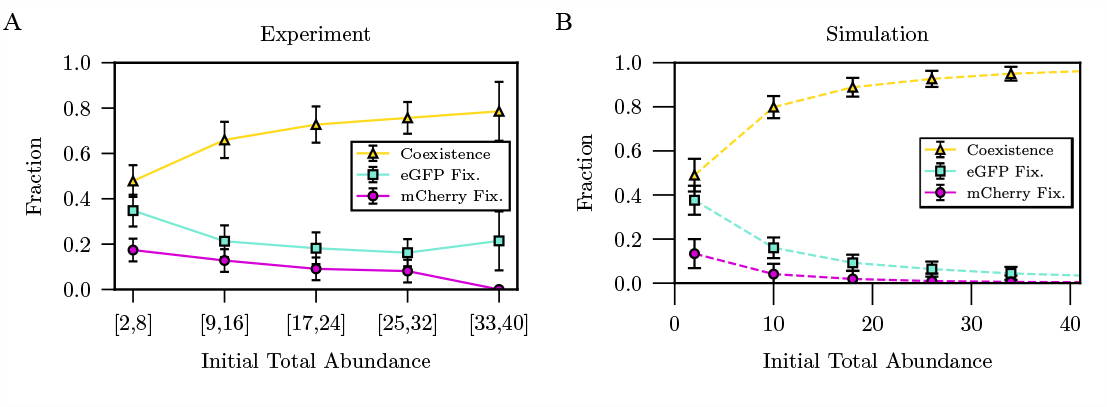
Likelihood of single-strain fixation or coexistence as a function of the number of cells seeded within the microchannels. A) Experimental results for five ranges of initial total cell abundance. Both cell types, under high induction, were randomly loaded into the microchannels at equal concentrations. Sample size from left to right: N = 69, 47, 33, 36, 14. B) Simulation results (*N* = 5000 for each data point) for an increasing, equal abundance of cells of each type randomly positioned within the chamber. Error bars result from bootstrapping the experimental data.

### Cell morphology has minimal influence on the ecological diversity

We further proceeded to test how differences in cell shape affect community fixation or coexistence. By exposing *E. coli* to the small molecule S-(3,4-Dichlorobenzyl) isothiourea (A22), which inhibits MreB (an actin-like cytoskeletal protein that participates in the maintenance of the rod-like (bacillus) morphology of *E. coli* [4, 48]) we are able to tune the aspect ratio of the cells to a rounder (coccus) morphology (see *SI Figs. S6, S12*). We find this shape change to have little effect on the doubling times of the cells both in bulk and in our microfluidic devices (see *SI Figs. S7, S8*) in agreement with previous reports [9]. We then performed a series of microchannel competition experiments, in the presence of A22, between *E. coli* (MC1400) cells and an A22-resistant *E. coli* mutant (E143A). The now more coccus strain was made distinguishable by mCherry expression while the mutant strain remained bacillus and expressed eGFP. At high L-arabinose induction levels, as presented here, the growth advantage of the green cells over the red cells, as observed in the original bacillus/bacillus experiments, was maintained. Microchannels were initially seeded at low abundance as described previously.

Figure 4A (*Movie S3, S4)*) outlines the outcomes of these experiments. While competition between the two bacillus strains resulted in the faster growing strain (eGFP) fixating almost twice as often as the slower one (mCherry), the mixed morphology experiments show the coccus cells either coexisting with or primarily out-competing the bacillus cells. This morphological difference appears to provide a selective advantage to the slower growing strain that more than accounts for its reduced doubling time. Surprisingly, however, agent-based simulations showed that varying the aspect ratio of the coccus species had little effect on the competitive outcomes (Fig. 4B). This insensitivity to morphology persisted in simulations for mixed competition between cells with aspect ratios well beyond what we could access experimentally.

**Figure 4.**
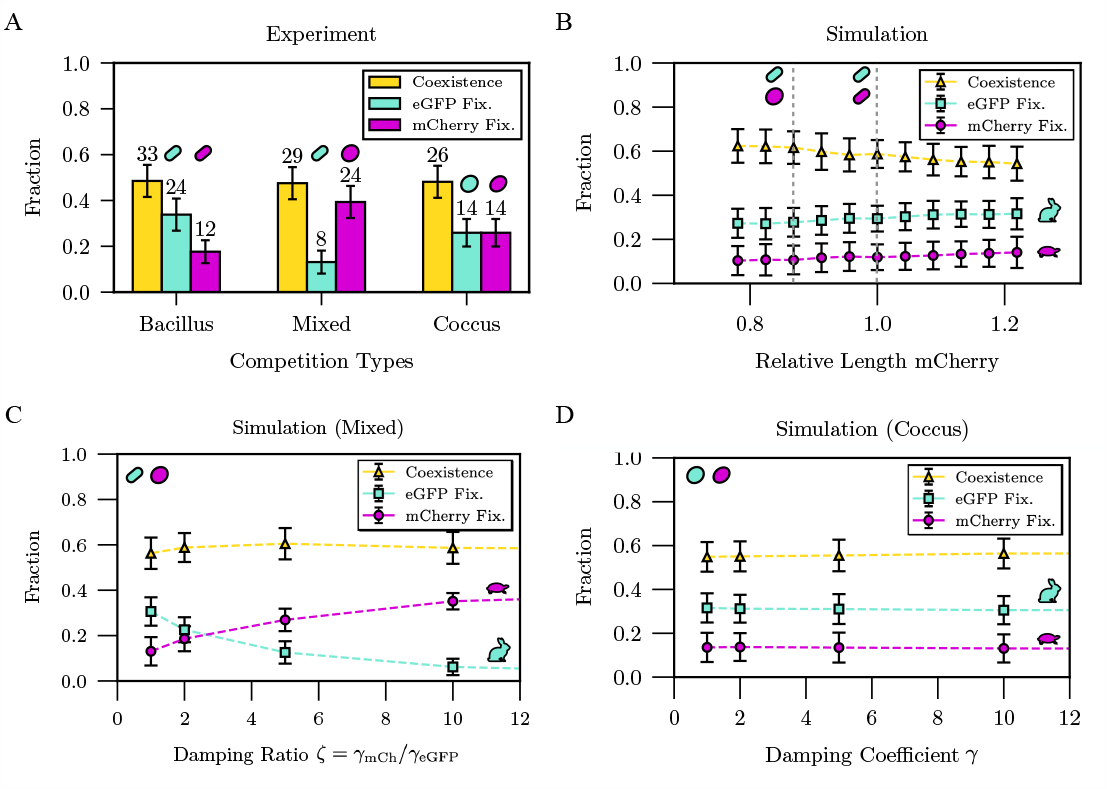
Fraction of competitive outcomes resulting in fixation or coexistence for cells with different morphologies (bacillus or coccus). All experiments/simulations were performed at high induction. A) Experimental outcomes in microchannels between two competing bacillus populations, a bacillus and coccus population (“mixed” competition), and two coccus populations. B) Simulated effects on outcomes within mixed competition by varying the relative length of the cells (length mCherry / length eGFP) while holding the eGFP cell length fixed (the grey dashed lines indicate the corresponding experiments), C) Simulations varying the damping coefficient of mCherry (coccus) cells relative to that of eGFP (bacillus) cells, and D) Simulations varying the damping coefficients of eGFP (coccus) and mCherry (coccus) cells in concert without changing their relative damping ratio. Faster (slower) growing cells are indicated by the rabbit (turtle). Error bars result from bootstrapping the data.

To account for our experimental observations, we hypothesized that the coccus cells may display significantly different hydrodynamic and surface adhesion properties than the bacillus cells. The wider coccus cells are expected to experience a higher frictional force from the top (PDMS) and the bottom (coverslip) of the ∼ 1 *μm* monolayer they are sandwiched between. To account for these factors, we systematically increased the damping for the coccus strain in our agent-based models. As shown in Fig. 4C, a high damping ratio (*γ*_*mCh*_*/γ*_*eGF P*_) does indeed favor the fixation of the coccus strain among the fixating trajectories while, once again, having little effect on the probability that the two strains reach coexistence.

We then performed a final set of competition experiments between two coccus strains (MC1400 cells expressing either eGFP or mCherry, at high induction, and in the presence of A22). Surprisingly, we observed effectively neutral competition between the two strains (Fig. 4A) despite the maintenance of a ∼ 10% growth advantage by the eGFP expressing strain. This observation cannot be explained by an increased level of damping affecting the coccus cells (Fig. 4D). So long as the relative damping is equal, the competitive outcome should be similar to that observed between the two bacillus strains. While we cannot conclusively explain this last observation, it is likely that the mechanical properties of the two coccus strains differ giving rise to a variation in their relative damping when squeezed within the microchannels, which will be explored in future work.

### Mechanical interactions inherently limit ecological diversity within confining microchannels

To understand the principles of population assembly, we next studied the strain diversity in populations that reach coexistence and assess how many neutral strains a microchannel can support. As shown in Fig. 1, after the initial filling period during which most community fixations happen, the population typically reaches a state with nematically ordered lanes, each composed of a single strain (eGFP or mCherry), that span the length of the microchannel. Our experiments on bacillus *E. coli* either display fixation, resulting in a single domain of one strain (i.e., adjacent lanes of one color/strain), or give rise to 2-4 domains of the two coexisting strains (Fig. 5A). While the dimensions of the microchannel can, in theory, support up to 12 single-lane domains, we never observed more than 4 either for the effectively neutral (low-induction) or the biased (high-induction) competition, This implies a significant, mechanically induced restriction to the ecological diversity of bacteria growing within these channels.

**Figure 5.**
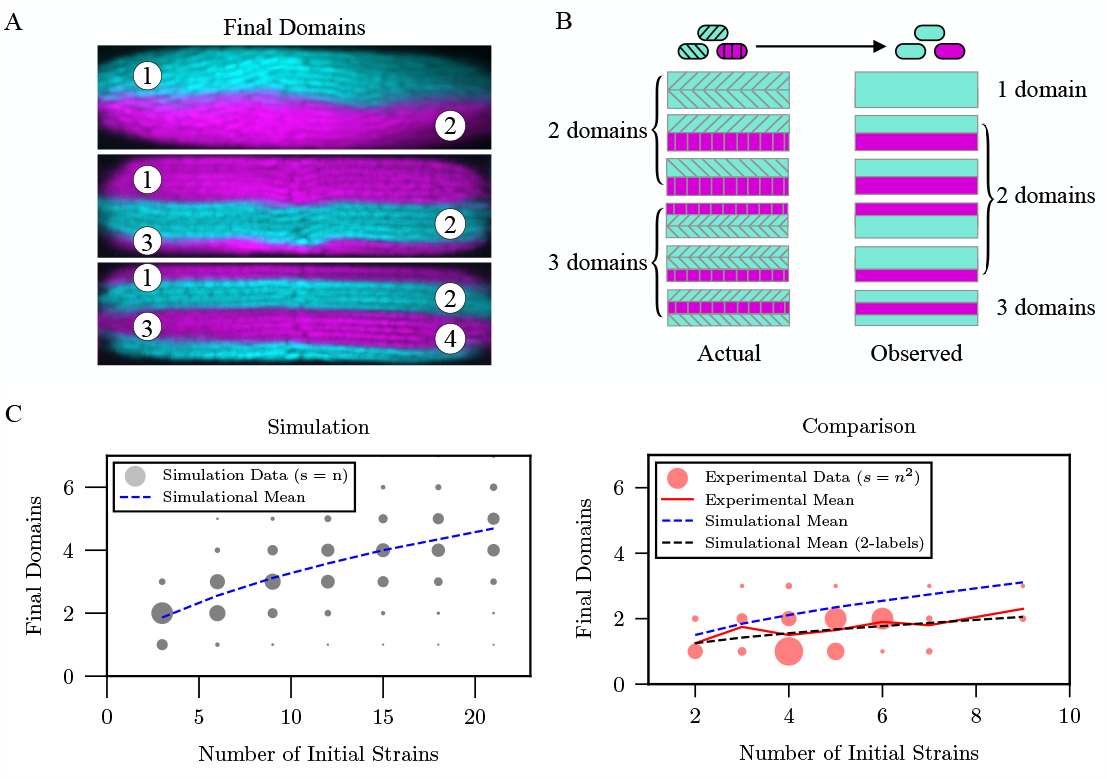
The diversity of coexisting populations is inherently limited by mechanical interactions within microchannels. A) Coexistence can accommodate a limited number of domains of each strain. B) Schematic example of the reduction in the number of observed domains when 3 strains (hashing) are indicated by only 2 labels (red/green). The observed number of domains will, on average, be less than than total number of strains that are actually present in the population. C) Simulation results for the number of surviving progenitor cell lines (domains) as a function of the abundance of seeded progenitor cells. The bubble plot represents the distribution of the data reflected by the bubble sizes (s = n). D) Comparison between experimental results and simulations. Both the actual as well as the dual-labeled simulation results (for comparison with experimental observations) are presented. The bubble plot represents the distribution of experimental data (bubble sizes, s = n ^2^).

Although our experiments employ only two bacterial strains, supported by simulations, they provide insight into a fundamental ecological question concerning the diversity of microbial communities supported by these microchannels. That is, if the channel were to be colonized by several different neutral strains, how many would be expected to remain in the coexisting state at long times? One way to address this question is to observe the dynamics of the clonal progenitor cells initially seeded into the channel assuming each to be an independent, neutral strain (irrespective of their fluorescent labeling). Due to the technical challenge of reliably tracking cell lineages within these microchannels over 24 hrs, we instead counted the number of red/green domains observed after 24 hours as a proxy. The number of domains provides a lower bound estimate for the number of surviving progenitor strains in the final population. This is because two adjacent domains resulting from progenitor cells expressing the same fluorescent protein will appear as one single domain, artificially reducing the number of observed domains (Fig. 5B).

On the other hand, simulations tracking the genealogy of the cells can directly identify the number of progenitor lineages that can be maintained in a microchannel. Simulations predict that the average number of surviving, neutral strains increases slowly with the initial number of progenitor cells, asymptotically converging to the maximum number of lanes (Fig. 5C). We can also use the simulations to retrieve the outcomes had we only two labels to compare with the experiments (Fig. 5B). The results of such an analysis are shown in Figs. 5C,D and agree well with our experimental observations. From these results, it appears that the final strain number grows sublinearly with the initial strain number–there is a diminishing return in the final diversity as the initial diversity increases.

### The diversity of dual-strain bacterial communities are well predicted by a simple Pólya urn model

Initial seeding conditions appear to determine whether competing populations will fixate or coexist, which ultimately determines the ecological diversity within these microenvironments. In this section, we show that a simple stochastic model known as a Pólya urn model [1,27,30] which does not explicitly incorporate any of the spatial or mechanical features of these systems, is largely able to predict the likelihood of strain fixation or coexistence, in good agreement with both our experiments and agent based simulations.

Pólya urn models are minimal models of constrained growth and history-dependent dynamics and have been applied to a range of phenomena – from the spread of infectious disease to the evolution of economic processes [19, 23, 41]. In Fig. 6A, we illustrate the Pólya urn model (see *SI Section 3*). The urn is seeded with two different elements at some initial abundance of each. At each iteration, an element in the urn is randomly chosen and returned to the urn along with an additional element of the same type. This process is meant to mimic the reproduction, growth, and preferential selection mechanisms observed in populations. The total population grows linearly in time providing a simple model of constrained growth. Through repeated iterations, the composition of the urn evolves, reflecting the changing population composition in time. The final state of any given run is a stochastic variable that strongly depends on the sequence of early events, and is drawn from a distribution determined by the initial strain abundance ratio [19, 23, 41].

**Figure 6.**
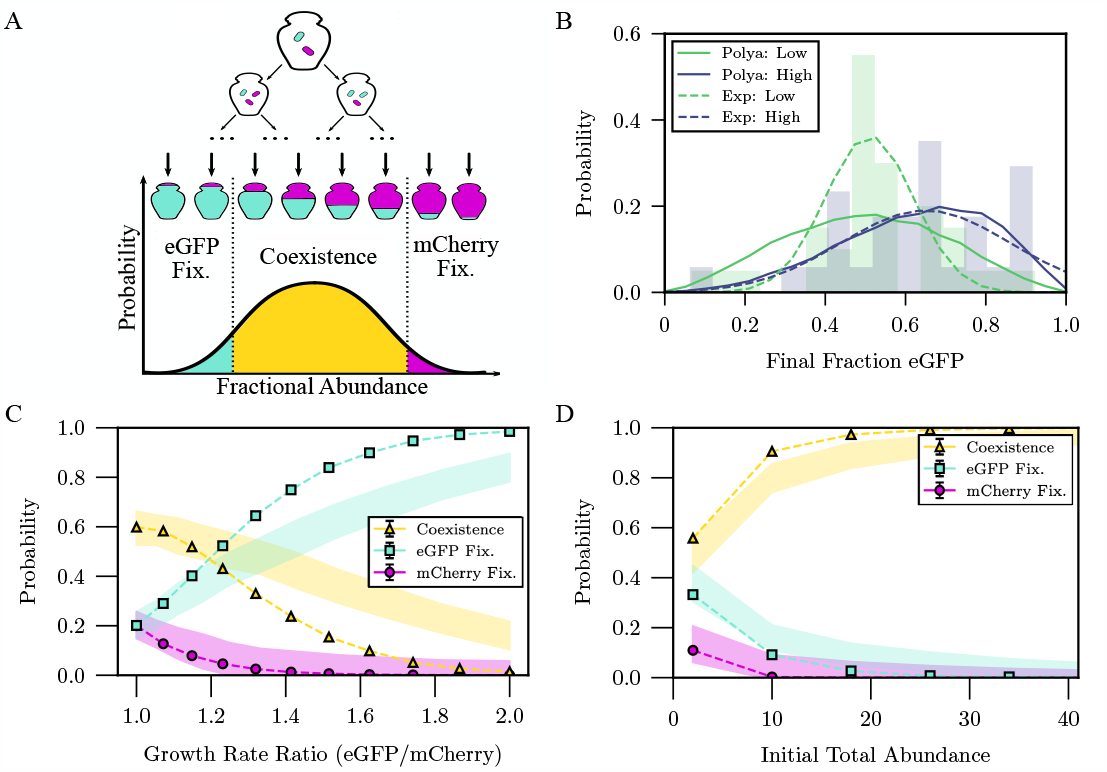
Coexistence and fixation in the modified Pólya urn model. A) Graphical representation of the Pólya urn process. Fixation of a strain is defined as occurring when the final fraction of the other strain is ≤ 0.2, otherwise the species are considered to coexist. Using this definition of fixation and coexistence, the competitive outcomes of the Pólya urn model semi-quantitatively recover the outcomes of agent based simulations (which were in agreement with our experiments). B) The probability of the fraction of eGFP cells in coexistence increases as the growth advantage of eGFP cells increases with the same trend in both experiments and the Pólya urn model (Pólya: High = 0.65±0.20; Pólya: Low =0.50±0.15; Exp: High = 0.65±0.21; Exp: Low = 0.51±0.11). The experimental means and standard deviations are from a Gaussian fit to the data, which approximates the,*B*-distribution predicted by the Pólya model. Experimental data is at low initial seeding with Polya seeded at *N* = 3/strain. C) The probability of each competitive outcome starting from a single cell of each strain as a function of the relative growth rate between strains (*eGFP/mCherry*). D) The probability of each competitive outcome as the initial abundance of each cell is increased (starting at equal abundance). The curves are the results of 20000 trajectories of the Pólya urn model. The shaded regions indicate one standard deviation from the mean of the agent based simulations.

Classical Polya urn models cannot describe true fixation because the progeny of all initial species remains present at any time albeit potentially at a very low abundance. To incorporate fixation in these models, we assume that a species is effectively absent from the system if its fractional abundance drops below a certain threshold value, as shown in Fig. 6A. We found that an empirically chosen threshold of 20% semi-quantitatively reproduces the competitive outcomes observed experimentally and in our agent based simulations. Figure 6B compares the resulting population abundance of each strain when the two strains reach coexistence. While the Pólya model predicts a wider distribution in final abundances than is observed, the peak of the distributions well agree for both neutral and biased competition. Likewise, we find that the model reproduces the relative probabilities of community fixation vs. coexistence for both neutral and biased competition (Fig. 6C) in agreement with observations (Fig. 2D). The dependence on the initial seeding abundance (Fig. 6D) exhibits the same trends as in Fig. 3–increasing the initial abundance results in more coexistence even if one strain displays a competitive advantage.

## Discussion

In this study, we focused on a model microbial community composed of two bacterial strains competing for space within a confined micro-environment. We investigated the determinants of the ecological dynamics that result in either coexistence or community fixation of the populations. Our work combined experimental and theoretical approaches to provide insight into how purely mechanical interactions both shape and constrain the ecological diversity of a bacterial community. Within these confined systems, factors such as the initial seeding, growth rate, and cellular morphology affect the ecological diversity in unexpected ways, highly distinct from the well known results in unconfined populations.

We observe that the establishment of population diversity within the microchannels is biphasic. Competition results in either one strain taking over the channel (fixation), typically on the timescale it takes for a seeded population to fill the microchannel, or both strains form into a relatively stable community (coexistence). Competing bacteria benefit from even a slight growth advantage (∼ 10%). A faster growth rate enables them to align into stable lanes across the channels quicker than their competitors [8, 40, 47], which facilitates their survival. The advantaged strain will more likely dominate the channel and, should one strain fixate, it will more likely be the strain with the growth advantage. While these effects of a growth rate advantage might be expected, we also observe, counterintuitively, that strains with a significant disparity in growth advantage commonly coexist in stable, spatially segregated communities.

The initial abundance of the cells plays a critical role in determining eventual fixation or strain coexistence. Coexistence is increasingly favored at an increasing initial abundance when both strains are uniformly seeded across the microchannel. We have assumed throughout that the microchannels are uniformly seeded, but our results also provide an indication of what might be observed for non-uniformly seeded populations. To coexist, each population needs to form at least one lane spanning the channel.

Within a well mixed, abundantly seeded system, cells typically first develop numerous small clusters that each grow radially outward until contacting either another cluster or the channel edge. When there are many clusters of a single strain present across the microchannel, the odds of these clusters merging into one or more lanes spanning the channel are high, which is why abundantly seeded systems predominantly achieve coexistence.

In cases of low abundance initial seeding, the system typically evolves into a few large clusters as the populations achieve the required density for ordering inside the microchannels. Note that this is similar to non-uniformly seeding the channel. When the expanding wavefronts of these clusters collide, the cluster closest to the center of the channel, with high probability, pushes all others out to quickly establish dominance. We conclude that both the abundance and degree of mixing within the initial seeded populations critically determine the resulting community diversity.

We also showed that cellular morphology alone does not appear to substantially influence the competitive outcomes we observe. One might expect the coccus cells to invade neighboring lanes of bacillus cells more often, since they are less constrained to remain within ordered lanes, but this behavior is not supported by simulations. Instead, morphological changes likely alter both the hydrodynamic drag and the friction coefficients between the bacterium and the confining surfaces. Since the bacteria are sandwiched between the coverslip and PDMS, we suspect that the coccus cells, whose diameter is larger than the bacillus, will experience more hydrodynamic drag and surface adhesion effects, and so are harder to push out of the chambers. Likewise, membrane properties such as surface roughness or elastic response may vary along with morphology, which would also contribute to frictional affects within the microchannels. Wang et al. showed that MreB, as the scaffold that rigidly links to the cell wall, can increase the mechanical stiffness of *E. coli* and *B. subtilis* [49]. Similarly, the circumferential rigidity (i.e., the effective stiffness in the radial direction [25]) of coccus *E. coli* due to A22 treatment was found to be weaker than in bacillus *E. coli* [4]. These all point to physical mechanisms for a difference in frictional properties between the coccus and bacillus cells in our experiments. However, to explain our observations of balanced competition between the two coccus strains requires an additional mechanism that would lead to adhesion properties that differ between the two A22 affected cell types. While even more speculative, ROS activity, which should be enhanced within the mCherry strain, might provide such a mechanism as it has been found to induce a stress response that can alter the membrane structure of bacteria [14, 46]. While frictional effects, formally incorporated as a damping ratio within our simulations, are able explain the changes in fixation probabilities we observe for various competitive morphologies, further experiments are needed to confirm this hypothesis.

Our results also enable us to obtain new insights into the age old problem of understanding how ecological diversity is established and maintained in closed ecosystems withs strong competition [17]. Here we ask the following: How many coexisting neutral ‘species’ can confined microchannels support? For our dual-labeled experiments of effectively neutral populations, we found a striking lack of variety in the final domain configurations compared to the possible maximum, hinting at a suppressed level of diversity. In conjunction with simulations that enable us to track lineages of an arbitrary number of neutral progenitor cells, we showed that the ecological diversity of these systems is indeed severely limited by mechanical effects alone. Simulations predict a slow growth in the final number of surviving strains relative to the number of initial strains, which indicates that, as more species are introduced, there is a rapidly diminishing probability that a specific strain will successfully coexist. While this is a common feature of population dynamics models [22, 29], it’s striking how constrained is the diversity of these systems.

Despite the complexity of the the system dynamics, many of the experimental observations and the agent based simulations are recapitulated by a simple minimalistic description based on a Pólya urn model. The defining property of the Pólya urn model is that initial imbalances and early fluctuations in the urn’s composition tend to get ‘frozen’ in a self-reinforcing mechanism that leads to a random but statistically predictable final system composition. Consequently, early colonization conditions largely determine the outcome of cellular competition within these spatially driven microsystems with a cascade of early advantages particularly amplified in smaller (or non-uniformly seeded) initial populations. On the other hand, for high initial seeding, the final diversity tends to depend less on the initial abundance because imbalances in population abundance are attenuated, as are the odds of one population being positioned less favorably than the other within the microchannel. These results indicate that Pólya urn models can present a paradigmatic framework for understanding global features of ecological diversity in confined spaces within a wide range of systems within the same ‘universality class’–despite neglecting the details of spatial complexity.

Mechanical interactions between cells and with their environmental confines have the potential to shape both the dynamics and diversity of bacterial communities. Within the engineered ecological environments studied here, we have seen that competing bacterial populations are able to, and preferentially tend to, coexist despite significant imbalances in the growth advantage of one species over the other. Likewise, mechanical effects can put severe limits on the diversity of species that can be supported within these microenvironments. Our work highlights how the details of early colonization conditions can largely determine the subsequent architecture and ecological diversity of a bacterial community. Further single-cell measurements within more ecologically realistic systems are needed to determine the relevance of our findings to true bacterial consortia.

## Methods

### Bacterial Strains and Plasmids

We employed *E.coli* strains MC1400 and E143A, which are both derivatives of wild-type *E.coli* K12 bacteria. The E143A strain [7, 33] was originally constructed by a MreB point mutation in the MG1655 chromosome (mreBE143A-msfGFPSW (kan)). As a result, the E143A strain is resistant to the MreB perturbing drug A22 so is able to preserve its rod-like shape without significant changes in growth rate. Plasmids pBAD-eGFP and pBAD-mCherry, used for distinguishing the two competitor strains, were constructed from a pBAD LIC cloning vector (8A) backbone with fluorescent reporters eGFP or mCherry inserted to site 2817, respectively (See *SI Section 1, Fig. S11*). All bacterial strains mentioned in this paper were chemically transformed with either the pBAD-eGFP or pBAD-mCherry plasmid.

### Microfluidics and Fabrication

The microfluidic device molds were created using standard i-line SU-8 photolithography at the Center for Research and Application in Fluidic Technologies (CRAFT) at the University of Toronto. Each PDMS casted microfluidic device consisted of an array of open-ended microchannels (Length × Width × Height = 46 × 12× 1 μm) and interspaced by 5 μm. Two much deeper flow trenches (Width × Height = 50 × 10 μm), perpendicular to the growth channels, supplied nutrients and carried away waste products/cells at the open ends of the microchannels. Details on the mold fabrication protocol as well as the soft lithography can be found in the *SI Section 1 and Fig. S1*.

### Sample Preparation

Details on the bulk sliding assays can be found in the *SI Section 1*. For the single cell assays, mCherry/eGFP *E. coli* strains were first inoculated from a single colony on the cell plates and grown separately in 5 mL of growth medium (LB broth + 100 μg/mL Ampicillin/Carbenicillin + 0.01 or 0.1% L-Arabinose) at 37 ^*o*^C, 250 rpm. Overnight cultures were regrown separately with 100x dilution in 5 mL of fresh growth medium or MreB-disturbing medium (growth medium + 0.5 μg/mL A22 compound) on the following day for 2 - 3 hours. The regrowth cultures were then mixed with a 1:1 ratio and diluted to a total OD of 0.1 in the supply medium (growth medium/MreB-perturbing medium + 1% BSA). The inner surface of the microfluidic device was passivated prior to cell loading by pumping the supply medium at 500 μL/hr from one of the entry ports for 5 minutes using a 5 mL syringe and a Chemyx syringe pump (Fusion 101A). The diluted cell mixture was pumped into the microfluidic device at 150 μL/hr from the entry ports using two 5 mL syringes until most of the microchannels were loaded with bacteria. After cell loading, the loading syringes were replaced by two clean 5 mL syringes filled with the supply medium which constantly replenished the medium inside the microfluidic device at a rate of 150 μL/hr for the 24-hour competition experiments. The waste was collected in two 50 mL Falcon tubes from the exit ports.

### Time-Lapse Video Microscopy

Imaging was performed on an inverted epifluorescence microscope (IX81, Olympus) equipped with a CCD camera (Hamamatsu) and a controllable MicroStage (Mad City Labs) with encoders, along with the appropriate GFP and RFP filter sets. Cells were observed using an Olympus 100 UPlanFLN 100x, 1.3 N.A. oil immersion objective. The open-source microscopy software toolkit *μ*Manager [13] was employed for automated imaging control. A home-built incubating chamber was mounted to the microstage in order to maintain the sample temperature at 32 ^*o*^C. This temperature was chosen, as opposed to 37 ^*o*^C, to reduce sample drift. Additional details can be found in the *SI Section 1*.

## Acknowledgments

T.M, F.H. and J.N.M. acknowledge support from an NSERC Discovery grant and an NSERC New Frontiers in Research Fund Exploration grant. A.Z. and J.R. acknowledge support from an NSERC Discovery Grant. We thank members of the Zilman and Milstein labs for feedback on this work. We also thank the Navarre Lab for contributing the MC1400 strain and the Morgenstein lab for the A22-resistant E143A mutant strain.

