## Supporting Information for "Mechanics limit ecological diversity and promote heterogeneity in confined bacterial communities"

##### This PDF file includes:

Supporting text
Figs. S1 to S12
Table S1
Legends for Movies S1 to S8
SI References

##### Other supporting materials for this manuscript include the following:

Movies S1 to S8

### Supporting Information Text

#### 1. Methods

##### Photolithography and Soft Lithography.

Details of the fabrication protocol involve standard i-line photolithography using two negative photoresists (SU-8 2000.5 and SU-8 2010) and a micropatterned generator (uPG501 Mask Writer) for laying features at different heights. Two photomasks were designed in AutoCAD, along with alignment markers, to generate a layered device with two different feature heights: 10 µm deep supply channels and 1 µm deep growth chambers. A micropattern generator (uPG501 Mask Writer) was used to print features on chromium-coated soda-lime photomasks with a 390 nm LED light source. Printed features on the photomasks were then etched out during the development process to create a shadow mask for the lithography.

In brief, a master mold wafer was created with SU-8 2000.5 and SU-8 2010 (MicroChem) epoxy-based photoresists that were spin-coated to the appropriate thickness (1µm and 10µm, respectively) sequentially. Directly after each spin-coat procedure, the wafer was soft-baked before UV light exposure. A manual mask aligner (EVG 620 Mask Aligner) was used to cross-link the desired features on the SU-8 coated wafer by exposure to UV light through the photomask. The treated wafer was baked again and developed with the SU-8 developer to remove the unexposed substrates. Heat-resistant tape was used to cover the alignment markers while spin-coating the second layer of SU-8 to align the multiple feature layers. The tape was then removed before UV exposure to reveal the alignment markers. Markers on the wafer were then aligned manually to the markers on the photomask with the mask aligner to overlay the variable height features. Finally, the wafer was hard-baked to strengthen the resulting master molds. The master mold was treated with trimethoxysilane (Sigma) for 15 min to prevent peeling/cracking of the mold prior to soft lithography.

Microfluidic devices (Fig. S1) were created by casting a degassed 10:1 mix of polydimethylsiloxane (PDMS) and curing agent (Sylgard 184 kit; Dow Corning) on the master silicon mold, followed by a 25 min curing cycle at 125°C. Each PDMS chip was gently peeled off the master mold and cut to the desired size with punched-out entry/exit ports. Both the chip and a glass coverslip (22 x 50 mm #1.5) were cleaned with ethanol, blown dry with clean compressed air, and then bonded together with a plasma cleaner (Harrick Plasma, PDC-32G). Each microfluidic device consisted of an array of open-ended microchannels (Length $\times$ Height = 46 $\times$ 1 µm) with 12 µwidths interspaced by 5 µm. Two much deeper flow trenches (Width $\times$ Height = 50 $\times$ 10 µm), perpendicular to the growth channels, supplied nutrients and carried away waste products/cells at the open ends of the microchannels.

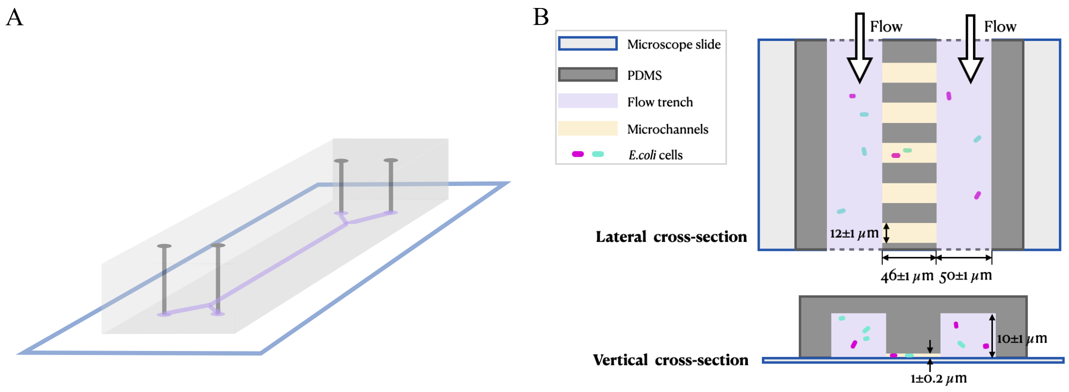


**Fig. S1.** Schematics of microfluidic device A) 3D structure of the microfluidic devices B) Detailed dimensions of the features on the microfluidic devices.

##### Sliding Assays.

The MC1400 *E. coli* strains harboring pBAD-eGFP or pBAD-mCherry, respectively, were first inoculated from single colonies on agar plates and grown separately in 5 mL of growth medium (Lysogeny broth (LB) Lennox + 100 µg/mL Ampicillin/Carbenicillin + either 0.01% or 0.2% L-Arabinose) at 37°C, 250 rpm. Overnight cultures were regrown separately at 100X dilution in 5 mL of fresh growth medium. Agar pads were prepared in 3.5 cm petri dishes with LB Agar (Miller) and the same concentrations of L-Arabinose and Ampicillin/Carbenicillin as the respective growth media of the cells. The regrown cell cultures were then diluted to an optical density (OD) of 0.25 and vortexed. One drop (1 µL) of the well-mixed culture was then deposited in the centre of each agar plate and allowed to absorb, covered, at room temperature for 30 minutes. The plate was then incubated at 37°C for 72 hours and imaged using an Invitrogen iBright FL1500 imaging system and analyzed with Fiji (ImageJ). The analysis consisted of splitting the red and green fluorescence channels and binarizing the signal. The "radial" areas of red and green domains occupied at a selected distance from the centre of the colonies were then measured, excluding the central region where the cells are mixed (Fig. S2). The green and red areas, as fractions of the total measured area, were then averaged across 3 samples and the mean and standard deviation were reported.

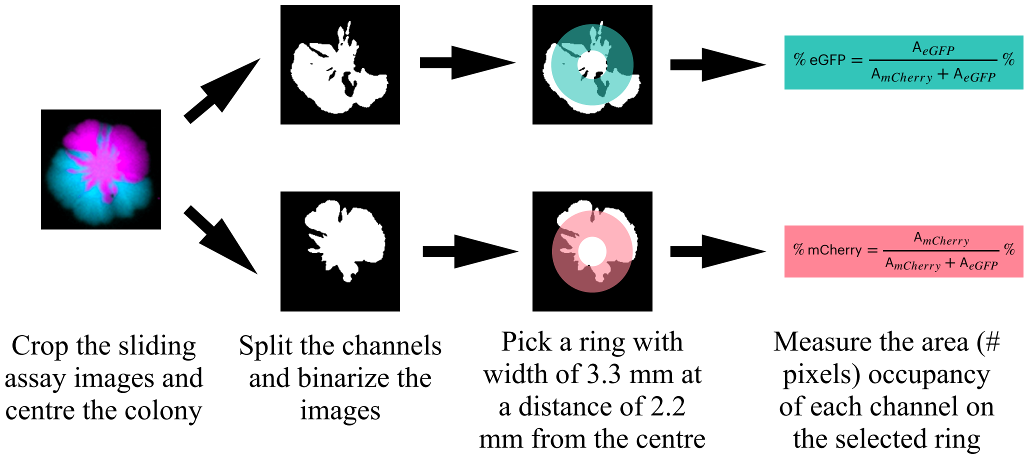


**Fig. S2.** Total fractional area occupancy analysis on the sliding assay. Fluorescent images taken from iBright imager were processed in µManager (contrast adjustment and thresholding). The pixel numbers on the binarized images at selected regions then were calculated in Python scripts as the area occupancy of each colony.

##### Time-Lapse Video Microscopy.

Imaging was performed on an inverted epifluorescence microscope (IX81, Olympus) equipped with a CCD camera (Hamamatsu) and a controllable MicroStage (Mad City Labs) with encoders, along with the appropriate GFP and RFP filter sets. Cells were observed using an Olympus UPlanFLN 100x, 1.3 N.A. oil immersion objective. The open-source microscopy software toolkit $\mu$Manager (1) was employed for automated imaging control. A home-built incubating chamber was mounted to the microstage in order to maintain the sample temperature at 32 °C. This temperature was chosen, as opposed to 37 °C, to reduce sample drift. A maximum of five parallel regions of interest, containing 3-5 neighboring growth chambers, were imaged concurrently and selected manually prior to each acquisition. Likewise, the focus was maintained through the software autofocus in $\mu$Manager. The exposure time was set to 125 ms for bright-field images and 300 ms for fluorescent images and the camera gain was set at a maximum (250). Bright-field images were taken every minute, while fluorescent images in the green and red channels were acquired every 11 minutes to minimize phototoxicity.

##### Single-Cell Analysis.

The fluorescent images were first processed in $\mu$Manager applying contrast adjustment, background subtraction, and image inversion for cell segmentation and tracking via DeLTA (2). The resulting single-cell information, such as cell dimensions and elongation rate, was then further analyzed with custom-built programs in Python to obtain the aspect ratios and doubling times (Fig. S3) under different conditions and their distributions among the populations. The doubling time is obtained by converting the elongation rate via:

$$T_d = \frac{\ln 2}{\alpha} \times \frac{60}{11} \quad [1]$$

where $\alpha$ is the elongation rate of individual cells from cell tracking in DelTA. The factor 60/11 is for converting the frame rate preset in Delta (seconds per frame) to our experiments (11 min per frame).

##### Error Propagation via Bootstraping.

To quantify the uncertainty in experimental measurements on the observed fraction of each competitive outcome, we bootstrapped 50 samples from the results (coexistence = a samples, green fixation = b samples, red fixation = c samples, a+b+c = sample size) 100 times with repeats. At each iteration, the new fractions of coexistence, green fixation, and red fixation were calculated. Then the histograms of these fractions from 100 iterations were fitted with normal distributions and the standard deviations of the distributions were taken as the error bars of the experiments.

The simulations were additionally used to predict the experimental error in observing each of the competitive outcomes. Note, the error bars shown for the simulations are not the error in the simulated predictions, which would be insignificant due to the large number of simulations we are able to perform (normally on the order of 1000s). Throughout, we perform bootstrapping in the agent based simulations by subsampling 50 data points. In our experiments, we regularly image between 50-100 microchannels for most data points, so we choose 50 subsamples for the simulation bootstrapping because it is on the order of the size of a reasonable experimental dataset. 50 subsamples was maintained as a standard throughout for all analyses of the simulations. This approach involves randomly selecting smaller groups of 50 data points from the much larger simulated dataset and calculating the standard deviation within these subgroups (Fig. S4). This process is designed to provide insights into the expected statistical characteristics of both the experiments we performed and provide a range for observations in further experiments.

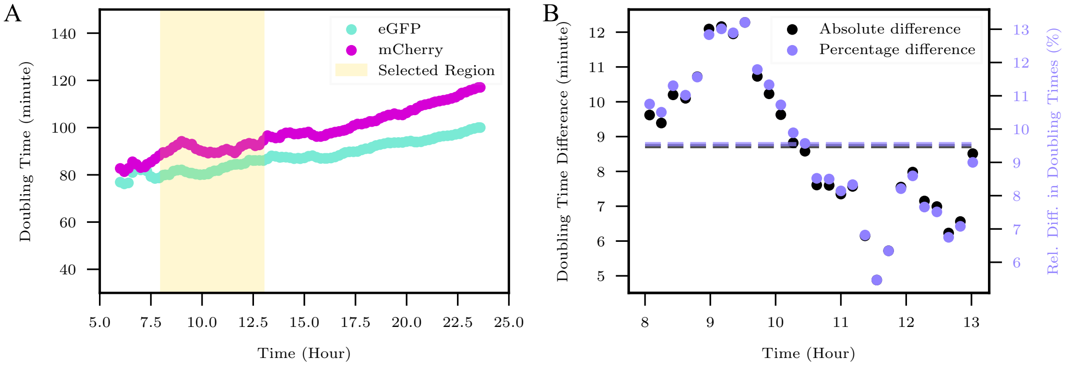


**Fig. S3.** Single-cell analysis on the bacterial doubling time of the eGFP and mCherry labelled cells in the competition experiments. A) The mean doubling time of individual cells at each frame for the competing strains is plotted as a function of acquisition time. The individual doubling time is converted from the cell elongation rate obtained from DelTA. The data from the initial colony expansion before the chamber filled (<5 hr in the example) are discarded due to insufficient cell tracking, as well as the "overpacked" state (>15 hr) where both strains grow slower. B) The absolute and percentage difference of the doubling time between mCherry and eGFP labelled cells within the selected window showed a stable growth advantage in the eGFP labelled cells.

#### 2. Agent based model

The Agent Based Model (ABM) is based on the models proposed by (3–5), which were successfully used to model spatially heterogeneous dynamics of bacterial populations in microfluidic devices. At the heart of these models lies the integration of cell-cell mechanical interactions, a critical factor in the bacterial population growth. In a virtual environment, individual bacterial agents act following rules governing their growth, division, and mechanical interactions with their neighbours and the environment. By including mechanical pushing among neighboring cells, the model captures the cooperative and competitive behaviours that shape bacterial colonies' spatial distribution and growth trajectories and enables observation of the populations abundance, providing interpretative lens for the experimental results. We used this approach to understand how the mechanical interactions among cells affected the diversity and the coexistence of different strains of cells in the open-channel microfluidic device.

In our ABM, cells are modelled in a 2D rectangular space, mimicking the environment of the manufactured microfluidic devices that contain a monolayer of bacteria. To mimic the experimental setup, the rectangular space has two parallel walls that do not allow cell crossings whereas the other two sides of the rectangle are open-ended (the long walls being of $44\mu m$ and the open ends measuring $12\mu m$). Similar to other studies of this nature, we have ignored directional flow of the medium in the channel, simplifying the dynamics of nutrient and cellular flow (4–7). Given the short width and depth of the channel, we assume an even distribution of nutrients necessary for growth and that no external flow moves cells. These assumptions are justified *post hoc* by the agreement with the experimental observations.

The bacterial cells are modeled as agents of spherocylindrical shape with average maximal length ($l_m$), width ($w$) and growth rate ($\alpha$). Each species $s$ is defined by these physical characteristics ($l_m^s$, $w^s$, and $\alpha^s$) that can be different between species. Each cell undergoes exponential growth along the long axis of the spherocylinder at a rate $\alpha^s$ keeping their width, $w^s$, constant during the entirety of their life-cycle (8–10). The orientation and center of mass of each cell, $\theta$ and $\vec{x} = (x, y)$ respectively, do not change unless acted upon by external forces.

The cells grow until they reach a maximal length, roughly two times their initial cell length, which is drawn from a uniform distribution on $[0.9\ l_m^s,\ 1.1\ l_m^s]$. Once they reach their maximal length, cells split into two daughter cells. Although the sum of the length of the daughter cells equals the length of the mother cell at division, the lengths of the daughter cells are not equal: the difference between the length of the daughter cells is drawn from a distribution ranging uniformly on $[0,\ 0.1\ l_m]$. The orientation and the center of mass of the daughters also differ from the mother such that the new orientation of the cells are $\theta + \delta_\theta$ and $\theta - \delta_\theta$, respectively, with $\delta_\theta$ drawn uniformly from $[-0.05\pi, 0.05\pi]$. The random draws of length and orientation aim to simulate the stochastic nature of cellular division processes. This randomness helps prevent division synchronization and strict nematic ordering within the microchannel.

As a cell grows and divides, it comes into contact with other cells. These mechanical interactions, along with the cell pushing up against the confining walls of the microchannel, are the source of external forces on the cell. Physical contact between cells occurs when the areas of two cells overlap, causing repulsion between the two cells. To depict the physical force arising from repulsion, we employ a soft sphere model of interaction for the portions of the cells that overlap. In this model, two overlapping soft spheres with widths $w_i$ and $w_j$ respectively are used, and each sphere is centered at the closest point between the central axes of two bacteria spherocylinders (11), as illustrated in Fig. S5. The position of the center of the sphere (i.e., the point along the axis of the spherocylinder closest to the point of contact) for cell $i$ is denoted as $\vec{r_i}$, and similarly, for cell $j$, it is denoted as $\vec{r_j}$, as depicted in Fig. S5.

The overlap of two soft spheres causes a repulsive Hertzian $\vec{F_{ij}}$ force along the normal direction $\vec{n_{ij}}$ between the two sphere

 Tianyi Ma, Jeremy Rothschild, Faisal Halabeya, Anton Zilman*, Joshua N. Milstein*

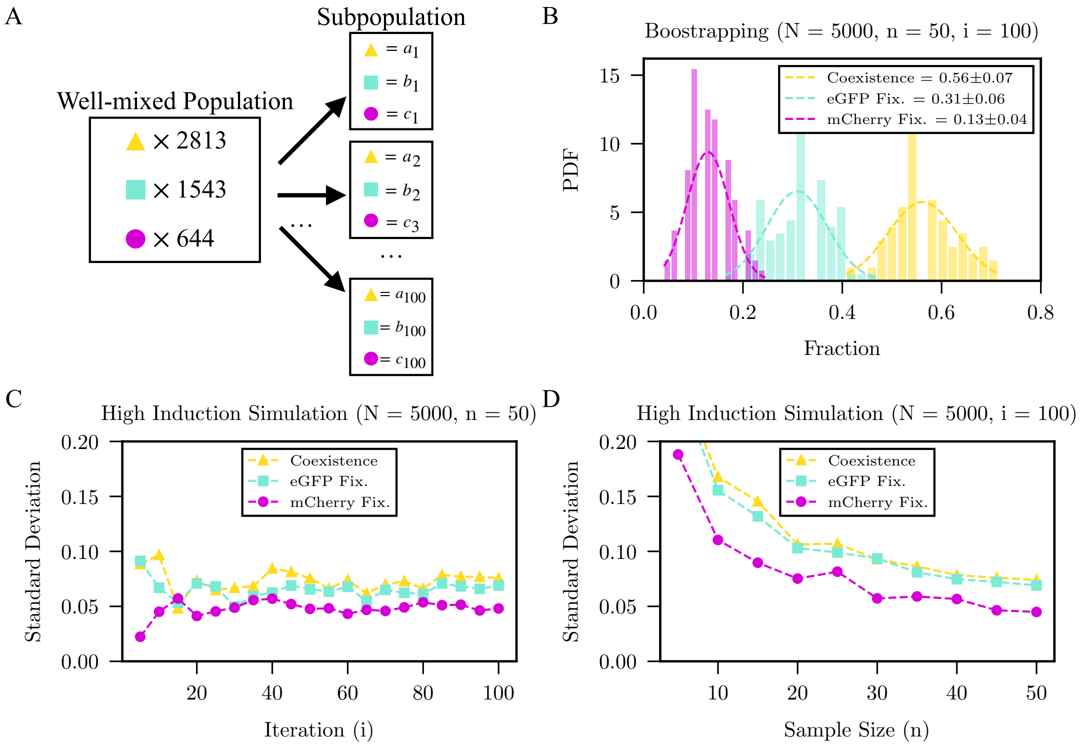


**Fig. S4.** Bootstrapping method A) Schematic of bootstrapping the simulation results. B) The distribution of the competition outcomes from bootstrapping 5000 simulations by subsampling 50 data points in 100 groups. The means and standard deviations are calculated from the Gaussian fits. The standard deviation from the bootstrapping converges quickly C) as the number of subsets increases (i.e., 50 subsamples randomly chosen and iterated $i$ times) and D) as the sample size increases (i.e., $n$ subsamples randomly chosen and iterated 100 times).

centers:

$$\vec{F}_{ij} = k_n \delta^{3/2} \vec{n}_{ij} \qquad [2]$$

where $k_n$ is a non-linear stiffness parameter related to the elasticity of the spheres and $\delta = (w_i + w_j) - |\vec{r_j} - \vec{r_i}|$ is the overlap distance between the two spheres (11). Note that $\vec{F}_{ij} = -\vec{F}_{ji}$. The center of mass and the orientation of cell $i$ change according to the following over-damped equations of motion:

$$\frac{d\vec{x}_i}{dt} = \frac{1}{m_i \xi} \vec{F}_i^W + \frac{1}{m_i \xi} \sum_j \vec{F}_{ij} \qquad [3]$$

$$\frac{d\theta_i}{dt}\hat{\theta} = \frac{12}{m_i \xi {l_i}^2}((\vec{r}_i - \vec{x}) \times \vec{F}_i^W) \cdot \hat{\theta} + \frac{12}{m_i \xi {l_i}^2} \sum_j ((\vec{r}_i - \vec{x}_i) \times \vec{F}_{ij}) \cdot \hat{\theta} \qquad [4]$$

where $l$ and $m$ are the length and mass of the cell, $\vec{F}_i^W$ is the force that is exerted on the cell from any contact with a wall, $\xi$ is a drag coefficient and $\hat{\theta}$ is the direction of the axis of rotation of the cell in the plane. The damping is proportional to the mass or inertia as in (4). The summation is over all cells $j$ that are in physical contact with cell $i$, otherwise no forces between the cells are generated. The force acted upon by the wall on the cell is similar to the Hertzian force in Eq 3 except that $\delta = w_i - |\vec{r}_i - \vec{d}_i^w|$, where $\vec{d}_i^w$ is the point on the wall closest to the cell.

Each cell persists in the simulation until its center exits the microchannel boundaries through either of the open ends. At this point, the cell is deemed to have exited the channel and is consequently eliminated from the simulation. This methodology enables the model to replicate the experimental dynamics, portraying cells navigating confined spaces, capturing their interactions via mechanical forces, and simulating their eventual departure from the channel.

Values for the numerical parameters used in these simulations are tabulated in Table S1, where the growth rate and maximal lengths were approximated from experimental measurements and the friction coefficient is has been sampled over the range of an order of magnitude. The remaining parameters cam be combined into one: $k_n/\rho\xi$, where $\rho = m_i/A_i$ is the density of a cell and $A_i = w_i((l_i - w_i) + w_i\pi/4)$ is the area of the cell. For our simulations, $k_n/\rho\xi = 1.25 \cdot 10^4 \mu m^{3/2} \mathrm{min}^{-1}$ which is an order of magnitude of (4).

In addition to the lateral confinement of the microchannel walls, the cells within the microchannel may come in contact with both the upper and lower boundaries (the coverslip and the base) between which they are sandwiched as a monolayer. The degree of this contact may depend on the bacterial shape and size. To account for this potential interaction, we took a

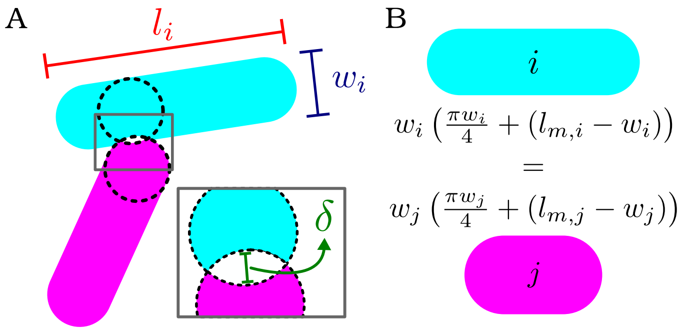


**Fig. S5.** A) Illustrative diagram of the contact between two cells and the overlap $\delta$ which is used to calculate the force. B) The $E.coli$ strains may have different geometries, however, their spherocylindrical shape and maximal area are conserved.

**Table S1. Simulation parameters for each figure in the main text.**

| Figure | Parameters | | | | | |
|---|---|---|---|---|---|---|
| | eGFP | | | mCherry | | |
| | $1/\alpha^{eGFP}$ (min) | $l_m^{eGFP}$ ($\mu m$) | $\gamma^{eGFP}$ | $1/\alpha^{mCherry}$ (min) | $l_m^{mCherry}$ ($\mu m$) | $\gamma^{mCherry}$ |
| 1F | 54 | 4.56 | 1 | 60 | 4.56 | 1 |
| 2D | 30-60 | 4.56 | 1 | 60 | 4.56 | 1 |
| 3B | 54 | 4.56 | 1 | 60 | 3.56 - 5.56 | 1 |
| 4B | 54 | 4.56 | 1 | 60 | 4.16 | 1 |
| 4C | 54 | 4.56 | 1 | 60 | 4.16 | 1-10 |
| 4D | 54 | 4.16 | 1-10 | 60 | 4.16 | 1-10 |

phenomenological approach and introduced rescaling of the damping coefficient to emulate the real-world phenomenon of cells potentially rubbing against the channel's surfaces. Through this adjustment, we aimed to enhance the model's accuracy in capturing the complex interplay between cell's changing shape and their interactions with the microenvironment. The new equations of motion with this damping coefficient are

$$\frac{d\vec{x_i}}{dt} = \frac{1}{m_i\gamma_i\xi}\vec{F_i}^W + \frac{1}{m_i\gamma\xi}\sum_j \vec{F_{ij}} \quad [5]$$

$$\frac{d\theta_i}{dt} = \frac{12}{m_i\gamma_i\xi {l_i}^3}((\vec{r_i} - \vec{x}) \times \vec{F_i}^W) + \frac{12}{m_i\gamma\xi {l_i}^3}\sum_j((\vec{r_i} - \vec{x_i}) \times \vec{F_{ij}}) \cdot \hat{\theta}. \quad [6]$$

All these equations are implemented in (12).

#### 3. Pólya Urn Model

Pólya urns are versatile probabilistic models that find applications across various fields including probability theory, statistics, and biology (13–15). These models serve as mathematical tools for understanding the dynamics of random processes involving sequential sampling and updating. This dynamic and self-evolving nature gives rise to interesting patterns and behaviours, making Pólya urn models classic models for exploring the interplay between randomness, competition and dynamics.

Starting with an urn occupied by an initial number $\alpha$ of red balls and $\beta$ blue balls, the evolution of the abundance of each type of coloured ball evolves according to the following iterative process. At each step, one of the balls is drawn uniformly at random from the urn. In the neutral Pólya urn model, the probability of drawing a ball of a certain colour, $p$, at each step is equal to the relative abundance of that colour in the urn at that time. The drawn ball is then returned to the urn along with an additional ball of the same colour before the next draw step. The abundances $n$ of red balls and $m$ of blue balls follows a stochastic trajectory, and the probability of finding $n$ red balls in the urn given a number of draws, $T$, is

$$P(n, T; \alpha, \beta) = \binom{T}{n-\alpha}\frac{\mathrm{B}(n, T-(n-\alpha)+\beta)}{\mathrm{B}(\alpha, \beta)} \quad [7]$$

where $\mathrm{B}(x, y) = \Gamma(x)\Gamma(y)/\Gamma(x+y)$ is the beta function (16). The count of total blue balls, $m = T - (n - \alpha) + \beta$, appears in the numerator of the equation. This equation is known as the Beta-binomial distribution, which can be extended into higher dimensions to the Dirichlet-multinomial distribution to simulate trials of the Pólya urn model with more than two colours of balls in the urn (17).

This Pólya urn model can be employed as a conceptual framework to study the dynamics of bacteria growing in a microchannel, offering insights into the evolution of the abundance of bacterial populations inhabiting the space over time. In this context, the microchannel serves as the "urn", while the bacteria represent the "objects" being drawn and reintroduced. As time progresses, bacteria in the microchannel will grow and divide. This can be modeled as a process of sequentially drawing a bacterium from the urn and then reintroducing a new bacterium into the urn where the newly introduced bacterium represents the offspring of the drawn bacterium. Just as in the Pólya urn model, attributes associated with the drawn objects (bacteria) can influence their reintroduction probabilities. In the context of bacterial growth, attributes include growth rate, fitness, or genetic characteristics. Bacteria with higher growth rates or better fitness might have a higher chance of being selected for division and reintroduction. As such, the model captures the stochastic nature of bacterial growth and division, which may be influenced by various factors such as nutrient availability, environmental conditions, and interactions between individual bacteria.

As stated above, fitness differences between bacterial strains, such as differing growth rates, can be incorporated into the Pólya urn model by including a fitness parameter to the probability of drawing an object from the urn. If a strain grows more rapidly, it is more likely to be selected for in a finite time step to give birth to progeny. In a 2 strain urn model, the probability for selection of the strain with abundance $n$ and relative fitness advantage $f$ is

$$p_n = \frac{f_n n}{f_n n + m} \tag{8}$$

where $m$ is the abundance of the other strains. Although an exact solution for the probability distribution of strains with fitness differences in the Pólya urn model is not available, we can simulate the system to find the probability of final abundances $P_f(n, T; \alpha, \beta, f_n)$ given initial abundances $\alpha$ and $\beta$ and fitness difference $f_n$, see code at (12).

Unlike the Pólya urn, which has an infinite carrying capacity, the microchannel can only contain a limited number of bacteria considering the physical limitations of the space. Nevertheless,the Pólya urn model's key characteristic is that it captures the probabilistic nature of sequential sampling and reintroduction, rather than any specific carrying capacity assumption, which turns out to be largely sufficient for our purposes. In the context of bacteria growing in a confined environment, the model's ability to simulate the dynamic interplay of growth and division while considering attributes such as growth rates, genetic diversity, and competitive advantage are the relevant features studied.

Another difference between the standard Pólya urn and our system is that the reintroduction scheme in the urn model does not allow for the permanent removal of any strains in the urn, unlike the extinction of strains which is observed in the microchannel. To relate the model to the growth of bacterial populations in a microchannel and describe the loss of diversity, we define a strain to be extinct when it's relative abundance falls below a threshold $\sigma$.In other words, although an abundance of species present initially never decreases to zero, if it's relative abundance becomes too low we consider it to be essentially removed from the microchannel as it is very unlikely to be selected for and continue providing individuals to the system. In the two strains case, the probability of extinction of a species is

$$P_{\text{ext.}} = \sum_{i}^{N\sigma} P_f(i, T; \alpha, \beta, f_n) \tag{9}$$

the probability of dominance is

$$P_{\text{dom.}} = \sum_{i=N(1-\sigma)}^{N} P_f(i, T; \alpha, \beta, f_n) \tag{10}$$

and the probability of coexistence is

$$P_{\text{coex.}} = 1 - P_{\text{dom.}} - P_{\text{ext.}} \tag{11}$$

where $N = n + m$. For the numerical simulations shown in the main text, we have chosen the number of draws/time steps to be $T = 2500$, which is a point where the distribution has converged numerically to a steady state probability of the relative abundance. To provide a more physical context to this choice, Within the context of the Polya Urn model, after a time $T = 2500$, the population in the system reaches a count of 2500 + the initial number of cells. In our experimental framework, initializing the system with 2 cells and allowing it to evolve for a duration of 12 hours yields an approximate population size of $2^{12} = 4096$. This count encompasses both cells present within the system and those that have exited the system, as the latter can be construed as undergoing growth and division albeit outside the confines of the chamber. Assuming that the total number of cells in both systems need to be similar, our choice of the Polya Urn model timescale roughly matches the experimental timescale, although further work is needed to more appropriately map the model onto the experiment. Testing various thresholds, we found that $\sigma = 1/5$ gave a surprisingly good fit with the quantitative experimental observation of the initial values of fixation and coexistence in the growth and density curves. However, more work is needed to determine the significance of the threshold value and understand its relation to the coexistence and fixation dynamics.

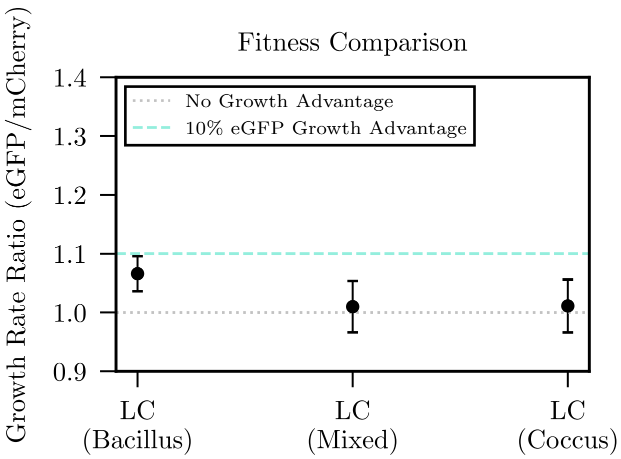


**Fig. S6.** Relative doubling times measured in liquid culture (LC), at high induction, between various strains employed. LC (Bacillus), LC (Mixed) and LC(Coccus) correspond to a comparison of the doubling times between: WT-mCherry (bacillus) and WT-eGFP (bacillus) in LB, WT-mCherry (coccus) and MT-eGFP (bacillus) in LB+A22, and WT-mCherry (coccus) and WT-eGFP (coccus) in LB+A22. Sample sizes are 12 replicates for all conditions.

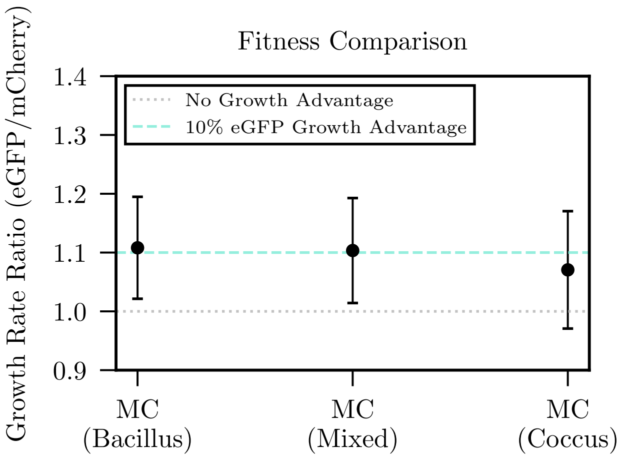


**Fig. S7.** Fitness difference of competing strains measured in the microchambers (MC) compared to liquid culture (LC) measurements (all at high induction level). The competing strains employed in different microfluidic experiments (Bacilli, Mixed, and Cocci) all display a growth advantage of 10% by the eGFP-labelled strains, when grown together, while the growth rate difference is about 6% in liquid culture (grown separately). Sample sizes from left to right are: 12, 6, 5

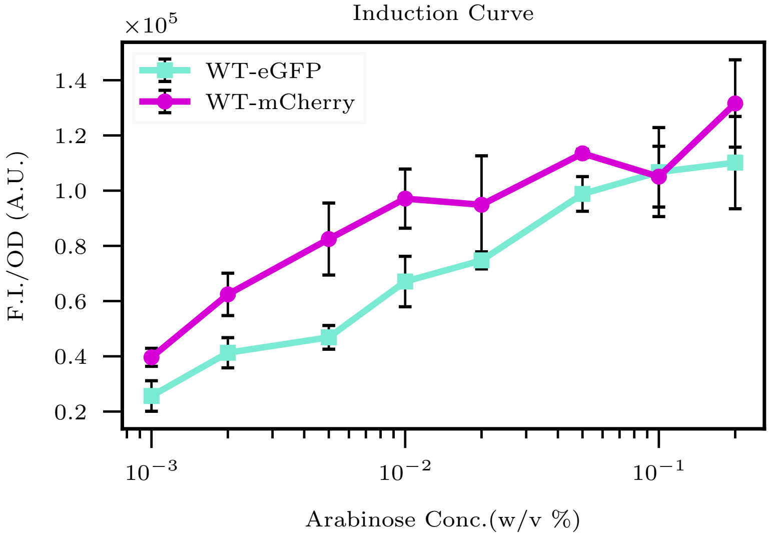


**Fig. S8.** Induction curve of the wild-type *E. coli* cells with eGFP and mCherry fluorescent reporters. The expression level of the population can be controlled by L-arabinose concentration. Low or high induction mentioned in the paper refers to a final concentration of 0.01% or 0.2% (w/v) L-arabinose in the growth medium, respectively. Error bars are the standard deviation of 6 replicates.

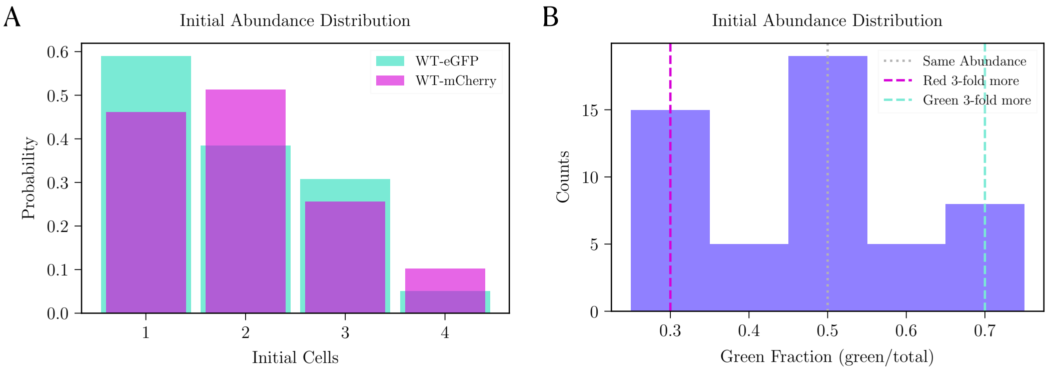


**Fig. S9.** Low abundance seeding conditions. Initial cell loading distribution for 'Bacilli', 'Mixed', and 'Cocci' experiments. Each strain has no more than 4 cells (A) and no more than 3 fold differences (B) between the competing strains.

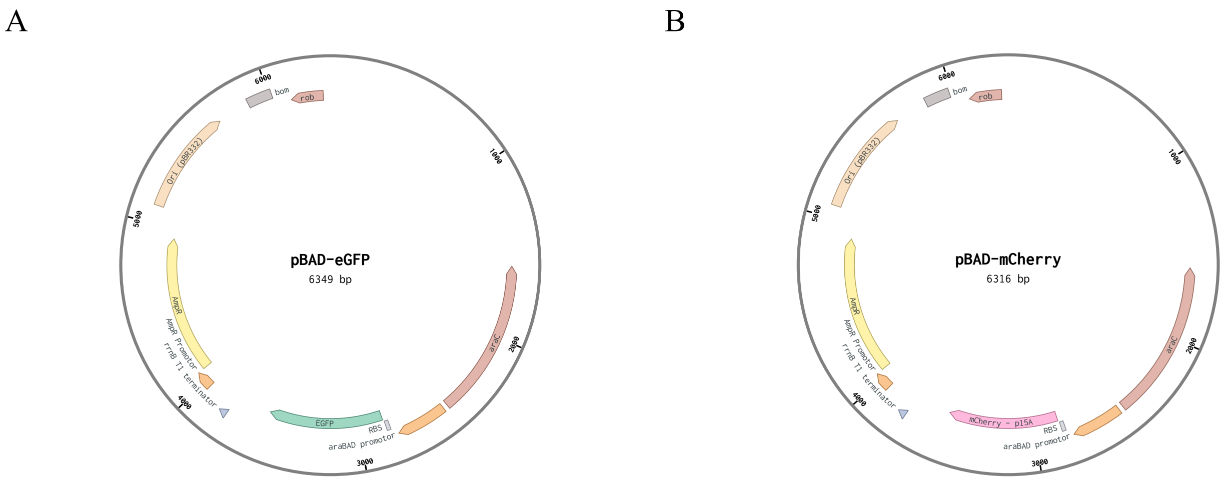


**Fig. S10.** Plasmid maps of pBAD-eGFP and pBAD-mCherry for distinguishing all the competing strains. eGFP and mCherry were inserted to site 2817 on the pBAD LIC cloning vector (8A) backbone.

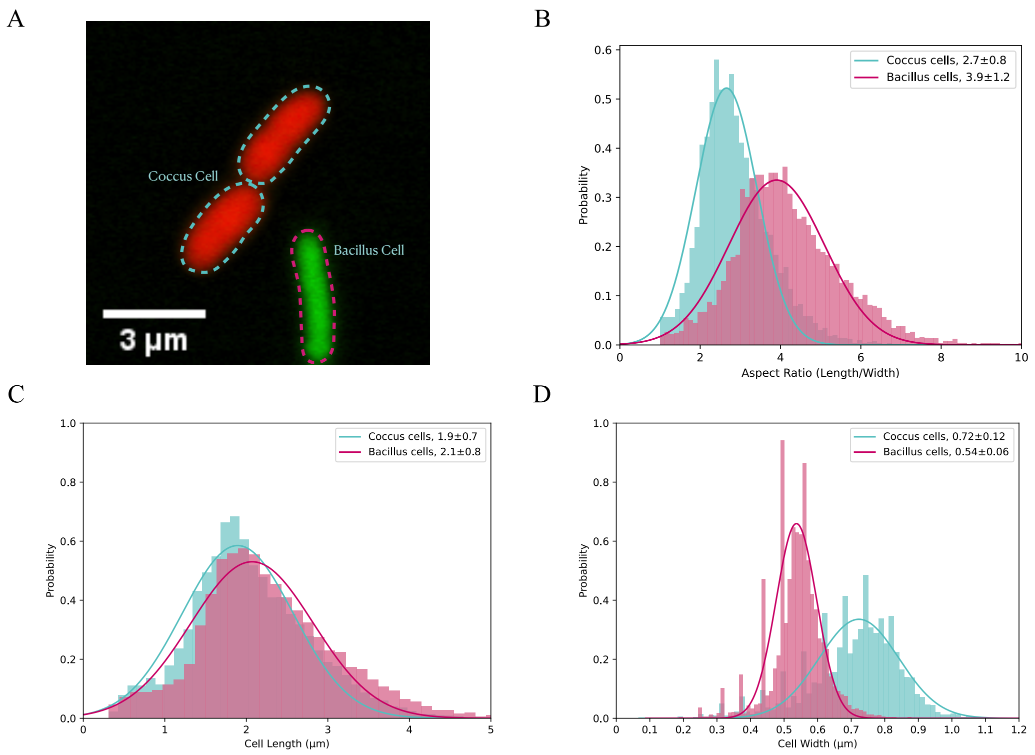


**Fig. S11.** Bacterial strains with different morphologies in the 'Mixed' competitions. A) A microscopic image of competing bacillus and coccus cells shows the difference in cell morphology. Distribution of B) aspect ratio, C) cell length, and D) cell width of the bacillus and coccus cells from single-cell segmentation fitted with a normal distribution.

 

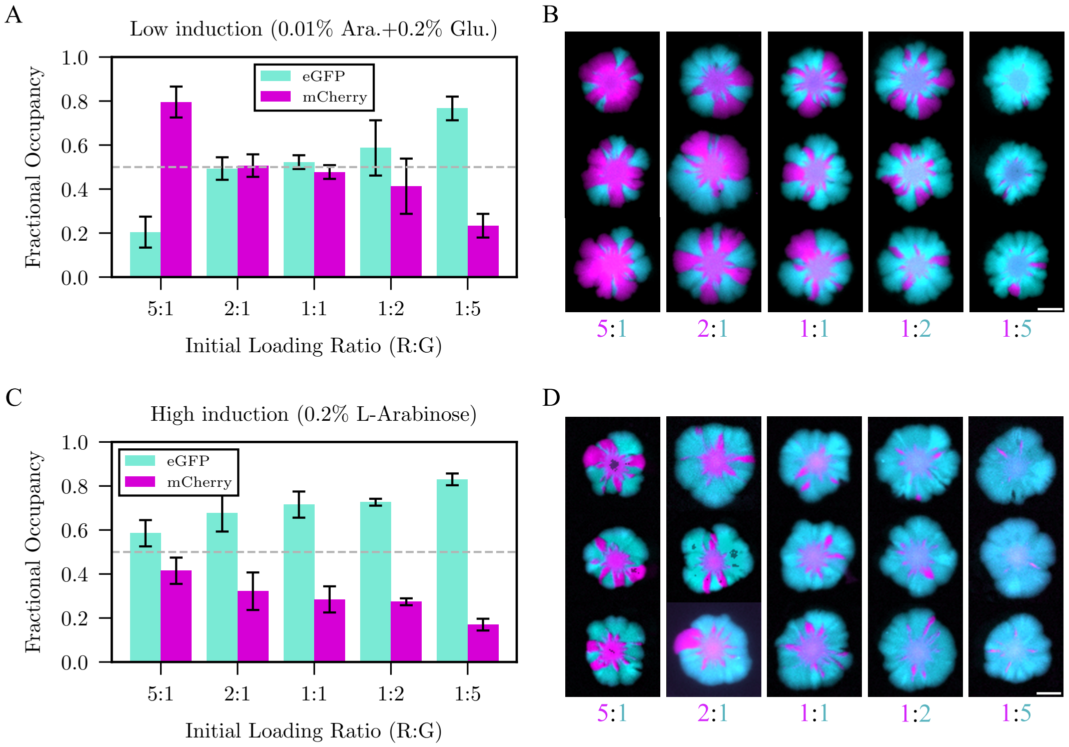


**Fig. S12.** Sliding assay and analysis of competing eGFP and mCherry labelled strains on agar pads with different initial $OD_{600}$ under low (A), (B) or high (C), (D) induction. (scale bars = 5 mm). The low induction condition was supplemented with glucose so that the colonies grew at a similar rate to those induced with additional arabinose. A) and C) Area fractional occupancy of each strain at a distance within 2.2 mm to 5.5 mm from the centre of the colony which corresponds to the sliding assay experiments shown in (B) and (D), respectively.

**Movie S1. Fixation Time-Lapse Video**

**Movie S2. Coexistence Time-Lapse Video**

**Movie S3. Mixed Time-Lapse Video**

**Movie S4. Cocci Time-Lapse Video**

**Movie S5. Fixation Simulation Video**

**Movie S6. Coexistence Simulation Video**

**Movie S7. Mixed Simulation Video**

**Movie S8. Cocci Simulation Video**
